# TEDEdb: a large-scale resource and multi-cohort analysis of transposable element differential expression in cancer

**DOI:** 10.64898/2026.04.27.721042

**Authors:** Gennaro Calendo, Morgan Chaunzwa, Iman Dehzangi, Jozef Madzo, Jean-Pierre J. Issa

## Abstract

The human genome consists of nearly 50% repetitive DNA, referred to for decades as “junk DNA”. These repetitive sequences, usually under the strict control of epigenetic silencing, have been observed to be aberrantly expressed in cancer. Some of these expressed sequences, e.g., transposable elements (TEs), can induce innate immune responses when de-repressed following treatment with epigenetic therapies. As a result, epigenetic therapy has been suggested to augment cancer therapies. TEs are traditionally ignored in most RNA-seq studies and their expression is often excluded from publicly available data sources. Thus, the vast amount of publicly available RNA-seq data is an untapped resource for exploring the role of TE expression in cancer and cancer treatment. Here, we present a uniform re-analysis of over 7,000 RNA-seq samples, encompassing more than 2,000 differential expression experiments across 220 cancer cell lines and 700 drug treatments. We observed that TE expression is more prone to batch effects than gene expression alone, necessitating the use of meta-analysis techniques to probe the dataset for global trends. We confirm that DNMTi and HDACis are powerful inducers of TEs. We also show that non-epigenetic compounds such as CDK and topoisomerase inhibitors can also induce robust up-regulation of transposable elements and confirm that this TE induction is consistent with viral mimicry response. We make all of the reprocessed data, web application, and database publicly available at: https://dataexplorer.coriell.org/TEDEdb/

## Introduction

The human genome is estimated to consist of roughly 54% of repetitive elements derived from mobile DNA [1]. Although most repetitive sequences consist of simple repeats in centromeric regions and telomere ends, currently active ones have been shown to contribute to structural variation [2] and regulate processes associated with transcription, chromatin accessibility, and inflammation response [3]. Activation of mobile transposable elements (TEs) can also induce various mutagenic processes, leading to cancer development and progression [4]. For example, autonomous long interspersed nuclear element-1 (LINE-1) retrotransposons self-propagate via RNA intermediates and have been suggested as biomarkers of neoplasia [5]. Moreover, hypomethylation during tumorigenesis can reactivate LINE-1 elements, resulting in disruptive insertions into cancer-associated genes such as APC in colon cancer and MYC in breast cancer [6–8].

TEs influence cancer progression through mechanisms beyond transposition into tumor suppressors or oncogenes. Genomic instability in cancer results in regions of DNA hypermethylation and hypomethylation [9, 10] and restructuring of chromatin [11]. Silencing DNA hypermethylation can occur in the promoters of tumor suppressor genes [12], whereas DNA hypomethylation is primarily observed in regions closely associated with repetitive DNA elements [9]. Activation of repetitive DNA elements can lead to a phenomenon known as viral mimicry [13, 14]. Viral mimicry represents a state of antiviral response triggered by endogenous signals that activates an innate immune response and upregulates interferon-stimulated genes [13, 15, 16]. This process increases sensitivity to cancer immunotherapies [17, 18]. By identifying cancers “primed” for viral mimicry through increased TE expression, clinicians may augment existing immunotherapy approaches or use TEs as biomarkers to stratify patients by their potential for therapeutic response.

Despite their clinical potential, TEs are often overlooked in transcriptome profiling, as typical RNA-seq pipelines struggle with the low mappability of highly repetitive, TE-derived reads [3]. This challenge arises because the repetitive nature of TE sequences leads to many ambiguously assigned ‘multi-mapping’ reads, requiring computational tools to accurately assign these reads while accounting for potential spurious assignments. To address this issue, several computational pipelines employ statistical assignment of reads [19–23]. Methods like TEtranscripts [19] and SalmonTE [22] typically provide counts at the TE subfamily level, losing locus-level specificity required to account for passive transcription. Tools such as Telescope [23], SQuiRE [20], and REdiscoverTE [21] offer locus-level quantification, with the latter being particularly attractive for large-scale studies due to its speed and accuracy, leveraging the Salmon [24] pseudoaligner and a novel index construction that avoids the computational cost of reassigning reads from precomputed BAM alignments.

Given the importance of TEs in cancer development and progression, along with recent advances in computational tools for their quantification, there is a pressing need and recent computational ability for large-scale studies to profile TE expression. Several large-scale pipelines exist for the uniform quantification of transcripts from publicly available sources. For example, ReCount [25] is one of the most comprehensive resources for uniformly processed RNA-seq data [26]. However, to our knowledge, no existing database contains quantifications specific to TEs at the locus level. For this reason, we developed a computational pipeline to quantify TE and gene expression from publicly available RNA-seq data. We employed this pipeline on over 7,000 publicly available samples across a variety of cancer cell lines and experimental treatments. Using this new resource, we confirmed the role of DNA methyltransferase inhibitors (DNMTis) as potent activators of TEs and inducers of innate immune responses. We also provide evidence to support recent findings implicating cyclin-dependent kinases (CDKs) as robust inducers of TEs [27–29] and viral mimicry activation [30]. Finally, to encourage further exploration, we created a database and a web-application called TEDEdb (Transposable Element Differential Expression Database) that are publicly available at: https://dataexplorer.coriell.org/TEDEdb/

## Results

### Re-analysis of transposable element differential expression from *>*2,000 experimental conditions

To examine global patterns of gene and TE dysregulation in cancer following treatment with epigenetic and/or cytotoxic drugs, we uniformly reprocessed over 7,000 publicly available RNA-seq datasets from SRA (see ‘Data Collection’) using a modified version of the REdiscoverTE [21] pipeline and performed annotation and differential expression analysis of more than 2,000 treatment vs. control contrasts (Figure 1A). These contrasts were composed of RNA-seq sequencing runs from 186 BioProjects containing 216 cell lines treated with more than 700 different compounds or experimental perturbations (Figure 1B). To ensure data quality, we estimated outlier status based on mapping statistics returned from Salmon [24] (see ‘Quantification’). BioSamples were flagged as potential outliers if their Median Absolute Deviation (MAD) exceeded 3 MADs for any one of the fragment-based statistics for paired-end and single-end sequencing runs, respectively (Supplemental Figure 1). Contrasts were flagged as outliers if any BioSample in the contrast was flagged as an outlier. Our analysis showed that most contrasts (1989 / 2215 = 90%) did not contain any BioSamples flagged as outliers. Importantly, all downstream analyses of TEs were performed on subfamily-level counts. This choice was made to simplify the interpretation of results in the context of the literature, which primarily reports TEs at the subfamily level. We would like to stress that locus-level quantifications are available in the database and were used to account for passive transcription.

**Figure 1.**
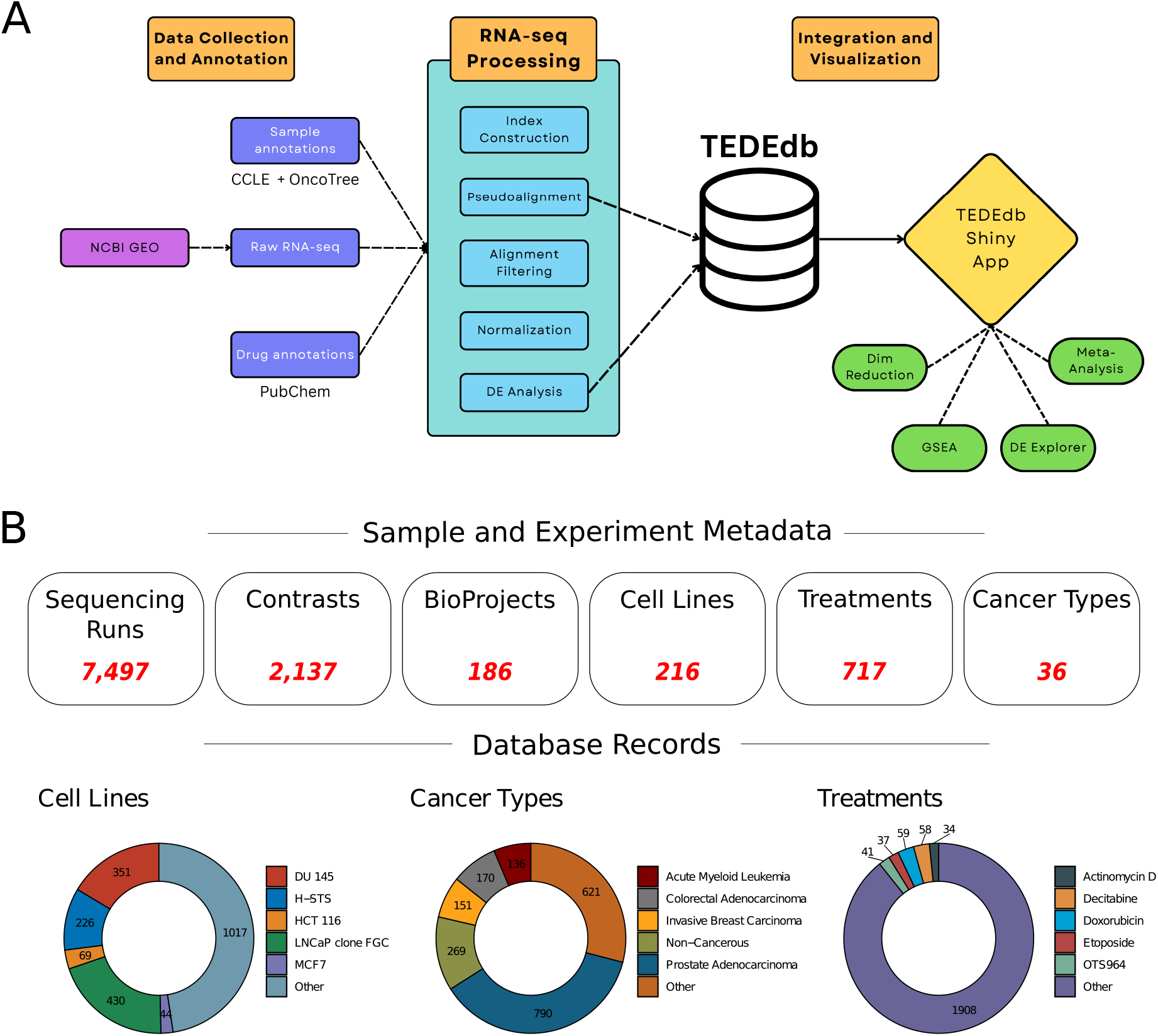
Overview of processing pipeline and database records. (A) Database Pipeline Overview. NCBI was queried for applicable studies. Following study selection, samples were uniformly annotated with information from Cellosaurus, The Cancer Cell Line Encyclopedia, PubChem, and Broad Drug Repurposing Hub. Raw fastq files were downloaded and uniformly processed using a modified version of the REdiscoverTE method for quantifying TE and transcript expression. Differential expression analysis was performed on the raw counts to create contrast-level data. Raw data and differential expression results were then deposited into the database. Differential expression-level data is then exposed to the user through the Shiny application. The application allows for interactive meta-analysis on selected differential expression data, dimensionality reduction, plotting of contrast-level volcano and MA plots, gene set enrichment, and over-representation analysis. (B) The database was constructed from 7,497 individual BioSamples and contains 2,137 records, representing the total number of treatment versus control comparisons for which differential expression testing was conducted and the results are available. The top row shows the unique number of metadata categories in the database. For each category, the five most frequent records are presented as donut charts below

### Transposable element expression is more prone to batch effects than gene expression

Batch effects are a well-known issue in high-throughput data analysis [31]. To illustrate the difficulty of combining sample-level count data from disparate experimental conditions and to examine gene and TE expression in response to technical factors, we examined the sample-level count data from control RNA-seq samples (i.e., RNA-seq sequencing libraries not treated with any compound) by gene and TE expression separately. After removing any samples flagged as outliers, TMM adjusted log2-count-per-million values for genes and TEs were used as input to principal component analysis. PCs representing ∼80% of the total variance were used as input for UMAP visualization of sample-level clustering. UMAP reduction on gene expression alone showed clustering by BioProject and cell line lineage (Figure 2). UMAP reduction on TE expression alone was more diffuse, with only certain cell line lineages, such as prostate and myeloid, showing larger clusters independent of BioProject of origin.

**Figure 2.**
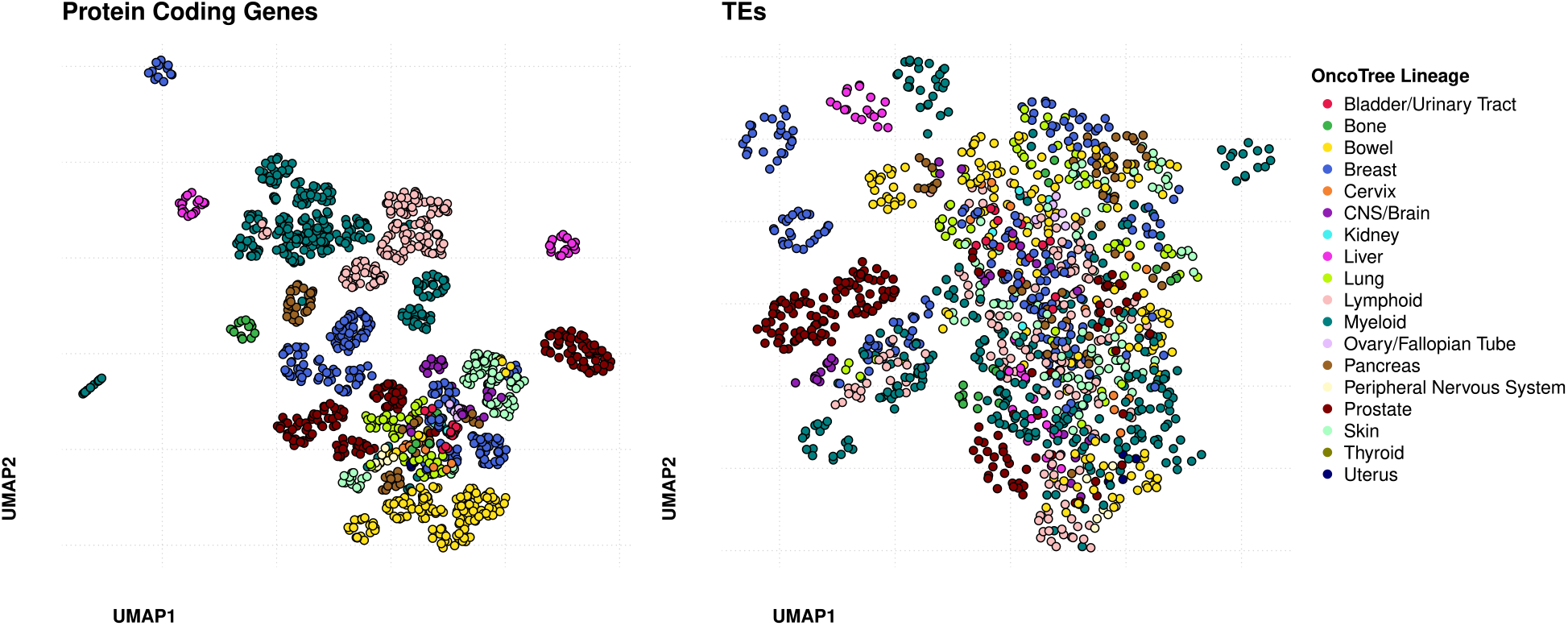
UMAP dimensionality reduction of BioSamples by gene and TE expression. (Left) UMAP dimensionality reduction on protein coding gene expression for control samples present in TEDEdb shows subclusters according to the primary lineage of the cell type. (Right) UMAP dimensionality reduction on TE subfamily-level expression alone. TE subclusters tend to be driven primarily by BioProject of origin rather than cell type lineage, indicating batch effects may be more pronounced for TEs than genes.

To quantify the susceptibility of gene and TE expression to technical noise, we performed sequential PERMANOVA, prioritizing cell lineage prior to the technical batch. Lineage explained only 19.5% of the variance in TE expression (*R*^2^ = 0.195, *p <* 0.001), while study origin captured the majority of the variance (*R*^2^ = 0.519, *p <* 0.001). Conversely, gene expression clustering was predominantly driven by cell lineage (*R*^2^ = 0.450, *p <* 0.001), with batch explaining a comparatively smaller fraction of the remaining variance (*R*^2^ = 0.374, *p <* 0.001). It is important to note, however, that cell lines and BioProjects are often confounded in the database (Supplemental Figure 2). Experimenters typically only assessed one or two cell lines of the same lineage in a given experiment, leaving batch and cell line information unable to be disentangled.

For these reasons, the primary focus of the database and downstream analyses is on differential expression results, where effect sizes can be compared and combined after considering the heterogeneity between studies. This approach assumes that while the absolute baseline of a feature may vary by lab, its transcriptional response to a stimulus is conserved. Such analyses are well-supported in the literature in the context of microarray meta-analyses [32–34], meta-analysis of RNA-seq [35], and TEs [36].

### Web application and database for the interactive exploration of results in TED-Edb

Since TEDEdb consists of over 2,000 differential expression results, we developed an easy-to-use web application to facilitate their interactive exploration. The application, located at:https://dataexplorer.coriell.org/TEDEdb/, consists of tabs that facilitate exploration and integration of differential expression results across studies. Figure 3 displays three analysis functions. The “Sample Selection” tab enables filtering of records by cell line information derived from Cellosaurus [37], the Cancer Cell Line Encyclopedia [38], and OncoTree [39] and compound information from PubChem and the Broad Drug Repurposing Hub [40]. Users can perform dimensionality reduction on the effect sizes of all genes and/or TEs from selected contrasts using the “Dimensionality Reduction” tab. In the example, all cell lines from Acute Myeloid Leukemia (AML) origins were selected for clustering based on the z-statistic of the differential expression. A small outgroup of contrasts treated with bromodomain inhibitors is observed, indicating a possibly conserved signature of response. Users can then select these contrasts and perform fixed-effect meta-analysis on the combined data (“Meta DE-Analysis” tab) to identify genes and TEs with evidence of conserved up- and down-regulation across the selected contrasts. In addition to these functions, the web application allows users to perform gene set enrichment analysis, over-representation testing, contrast ranking, and enables the exploration of individual experiment-level volcano and MA plots.

**Figure 3.**
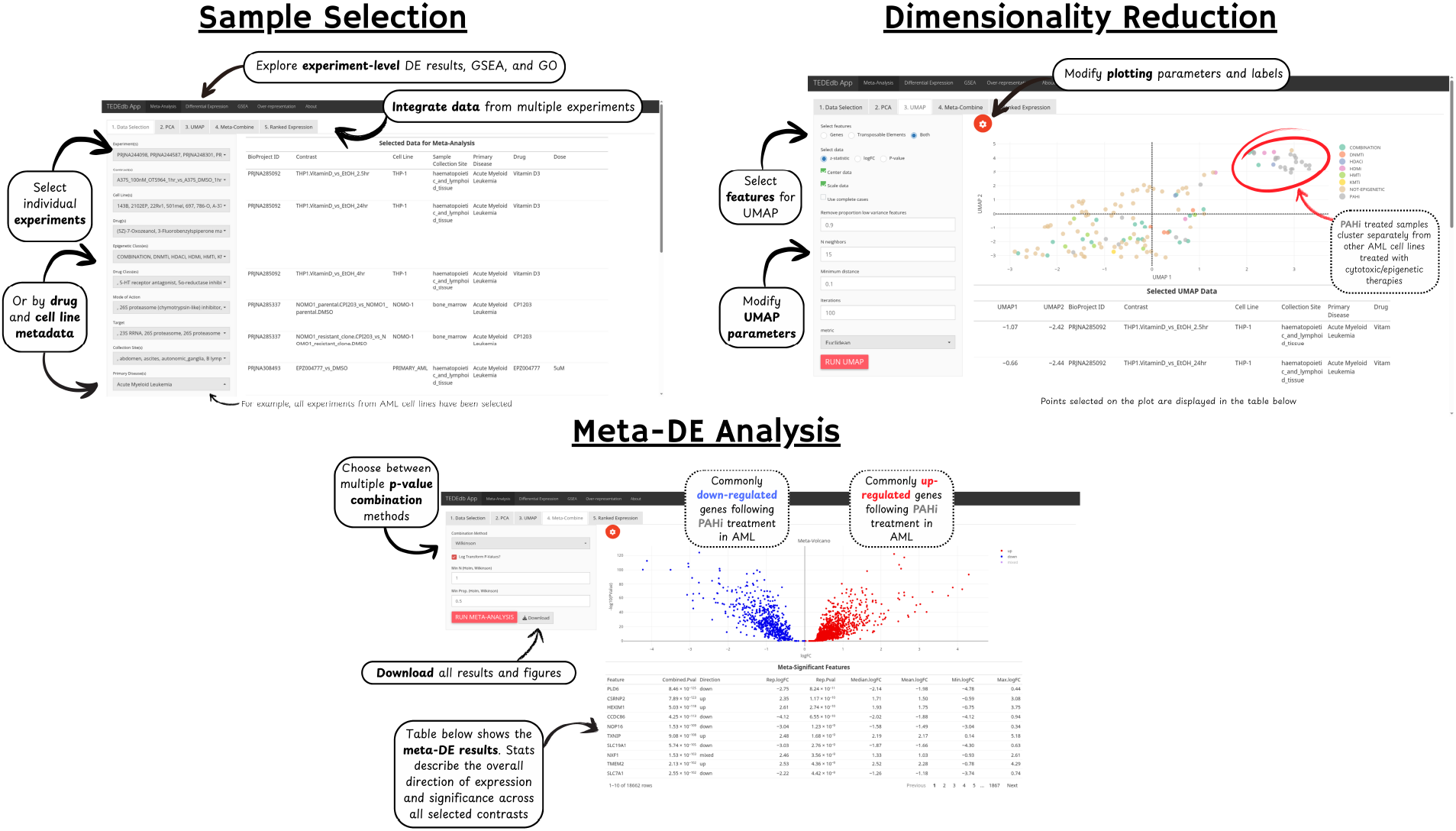
Overview of web application utilities. The Shiny web application allows for exploration of both contrast-level data as well as meta-analysis on collections of selected contrasts. A sample meta-analysis workflow is shown above. Top left: the sample selection screen allows users to combine metadata identifiers to be used in downstream analysis. In this example, all contrasts from Acute Myeloid Leukemia cell lines have been selected for downstream analysis. Top right: UMAP is performed on the feature-level z-statistic for all selected contrasts. The highlighted red circle shows that samples treated with proteins acting on acetylated histones (PAHi) show globally similar responses in AML cell lines. Bottom: p-value combination or fixed-effects meta-analysis can be used to combine differential expression results across studies. The meta-volcano plot shows genes and TEs that are both commonly up- and down-regulated following PAHi treatment in AML cell lines.

### Global analysis of TE differential expression reveals activation of TEs from non-epigenetic compounds

To describe the extent of gene and TE dysregulation in our database, we first examined summaries of differential expression results. We selected contrasts performed on cell lines tested with single compounds, excluding any contrasts containing BioSamples flagged as outliers, and extracted the percentage gene and TE dysregulation from each. We observed that in the majority of these experiments (820 / 1490 = 55%), less than ∼55% protein coding genes and even fewer TEs were dysregulated. However, a minority of contrasts (N=156) showed greater than 10% of TE subfamilies induced (Supplemental Figure 3).

To examine which classes of compounds resulted in this marked up-regulation of TE subfamilies, we examined average gene and TE percentage differential expression by epigenetic class. Histone deacetylase inhibitors (HDACi) and proteins binding to acetylated histones (PAHi) (e.g. bromodomain inhibitors) exhibited the largest range of protein coding gene dysregulation, 11.7 *±* SE 1.5% up- and 11.3 *±* 1.5% down-regulation and 12.6 *±* 1.6% up- and 16.1 *±* 1.5% down-regulation, respectively, with roughly the same degree of TE induction, 7.7 *±* 1.7% and 8.1 *±* 1.6%, as the other epigenetic classes, such as DNA methyltransferase inhibitors (5.1 *±* 2.1% up- and 0.7 *±* 0.6% down-regulated TEs) and histone methyltransferases (5.8 *±* 2.4% up- and 0.6 *±* 0.3% down-regulated TEs) (Figure 4A). Surprisingly, non-epigenetic compounds also induced TEs, with 3.6 *±* 0.3% average induction and 1.9 *±* 0.2% down-regulation.

**Figure 4.**
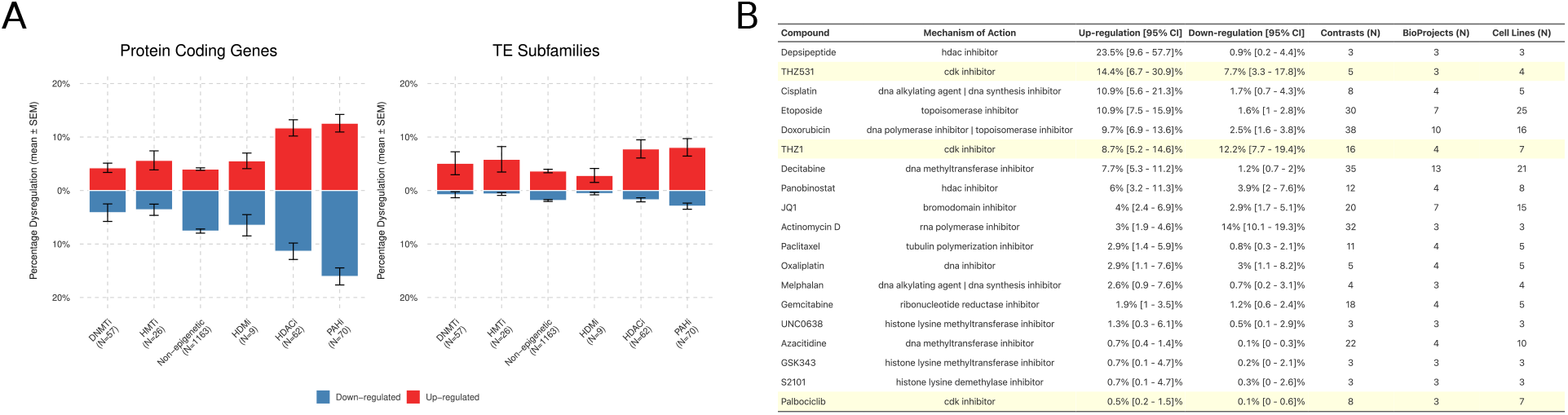
Average percentage dysregulated features by epigenetic class and mechanism. (A) Average +/- SEM percentage of dysregulated protein-coding genes for all contrasts within the given epigenetic class. (Right) Average percentage dysregulated TE subfamilies within the given epigenetic class. DNA methyltransferase inhibitor (DNMTi), histone deacetylase inhibitor (HDACi), histone demethylases (HDMi), histone methyltransferases (HMTi), proteins binding to acetylated histones (PAHi). (B) Table showing the mean of the percentage up-regulation and down-regulation [95% CI] of compounds present in more than two BioProjects. Contrasts (N); the number of individual contrasts averaged, Cell lines (N); number of unique cell lines across contrasts, BioProjects (N); number of unique BioProjects across all contrasts.

We then examined which compounds had the highest average percentage TE dysregulation. Figure 4B displays compounds ranked by their mean percentage TE up-regulation. The top-ranked non-epigenetic compound was the CDK12/13 inhibitor THZ531, which showed an average of 14.4% 95% CI [6.7%, 30.9%] up-regulation of TE subfamilies. THZ1, a CDK7 inhibitor, up-regulated an average of 8.7% 95% CI [5.2%, 14.6%] of TEs, similar to that of known TE inducing compounds such as Decitabine (7.7% 95% CI [5.3%, 11.2%]) (41). These results support previous studies suggesting the epigenetic role of CDKis towards the induction of silenced genes and TE expression [28, 29]. Genotoxic topoisomerase inhibitors Etoposide and Doxorubicin also resulted in up-regulation of TEs, with an average of 10.9% 95% CI [7.9%, 15.9%], and 9.7% 95% CI [6.9%, 13.6%] induction, respectively, both falling within the 95% CI of TE up-regulation by Decitabine.

To probe for shared patterns of gene and TE dysregulation across classes, we calculated the frequency, direction, and strength of differential expression across contrasts for all genes and TE subfamilies. Figure 5 shows the top ten genes and TE subfamilies in each class by their frequency of differential expression (adj.P.val *<* 0.1). We observed consistent up-regulation of LTR12C, LTR12D, HERV, and L1 subfamilies across the DMNTi contrasts, consistent with previous reports of viral mimicry induction [14, 41]. We did not observe many TE subfamilies or genes shared in the top sets across epigenetic classes. The top genes by CDKis were primarily down-regulated. Interestingly, CDKis showed consistent up-regulation of SINE Alu elements, unlike LTR and LINEs observed in other classes. It is important to note that CDKis in this context included multiple compounds with different cyclin-dependent kinase targets such as THZ1 (CDK7), Palbociclib (CDK4/6), THZ531 (CDK12/13) and MC180295 (CDK9/12).

**Figure 5.**
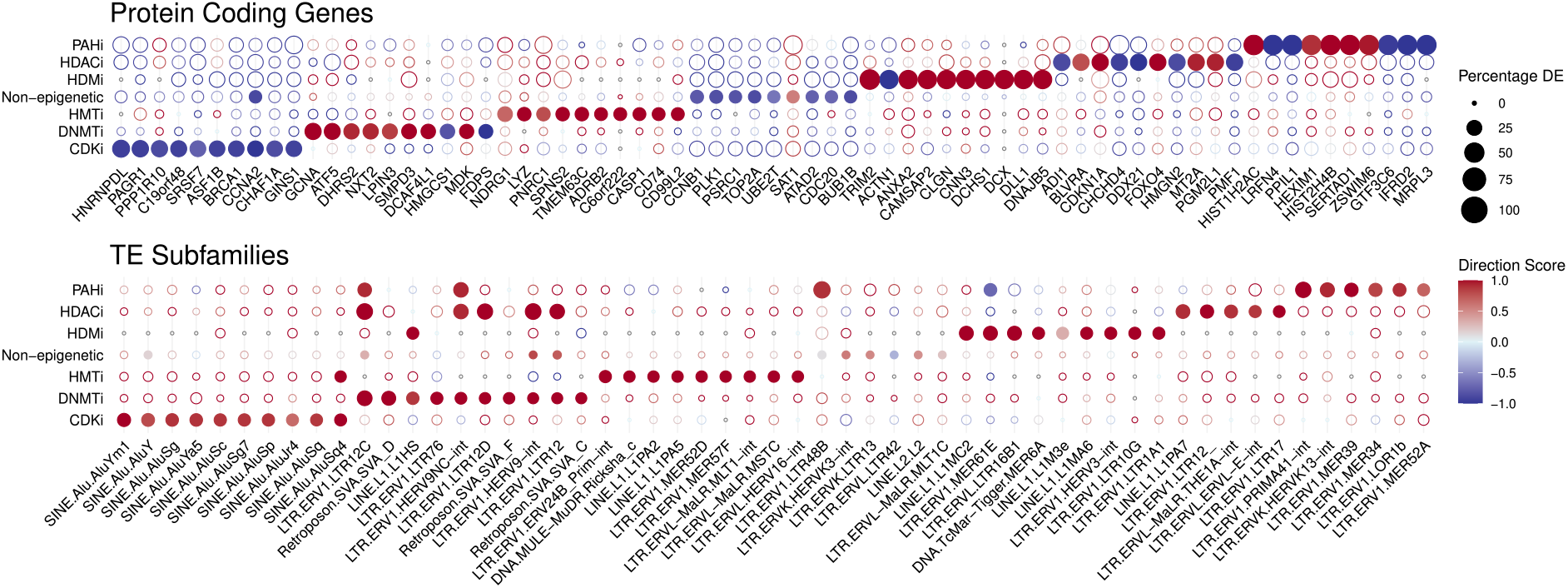
Dotplot of top dysregulated protein coding genes and TE subfamilies within each epigenetic class. (Top) protein coding genes. (Bottom) TE subfamilies. Filled circles represent top 10 features within that class by percentage dysregulation. Direction score was computed as (N up-regulated - N down-regulated) / total dysregulated. Positive scores indicate dominant up-regulation, negative scores indicate dominant down-regulation. Features were defined as differentially expressed at FDR *<* 0.1. Percentage DE represents the percentage of contrasts in which a particular feature was determined differentially expressed.

Finally, we sought to examine the global relationship between TE induction and interferon response and inflammation gene signature enrichment. We categorized all contrasts by their percentage of up-regulated TEs into tertiles and their induction of viral mimicry pathway responses (Supplemental Figure 4) as determined by competitive gene-set testing. Across all pathways we observed a significant increasing trend of enrichment of viral mimicry signatures with increasing induction of TEs (IFN alpha: *χ*^2^ = 27.161, df = 1, p-value = 1.872 *×* 10^−7^; IFN gamma: *χ*^2^ = 17.131, df = 1, p-value = 3.489 *×* 10^−5^, Inflammation response: *χ*^2^ = 47.295, df = 1, p-value = 6.105 *×* 10^−12^) consistent with the viral mimicry hypothesis. However, we also acknowledge the possibility of confounding by genotoxic treatments which may cause similar TE dysregulation and pathway responses.

### Meta-analysis yields a conserved signature of viral mimicry response across epigenetic and non-epigenetic treatments

Observations from above suggested that compounds traditionally considered non-epigenetic resulted in signatures of viral mimicry similar to those of canonical epigenetic compounds. In contrast to the descriptive analysis above, to generate specific signatures of conserved gene and TE expression, we performed fixed effects meta-analysis across contrasts from selected compounds. To select compounds, we first probed our database for the top compounds tested in multiple experiments, requiring those that were tested across the greatest number of BioProjects with the highest diversity of oncogenic lineages. The top compounds fitting this description were, Decitabine (Contrasts: 35; BioProjects: 13; Cell Lines: 21; Lineages: 7), Doxorubicin (Contrasts: 38; BioProjects: 10; Cell Lines: 16; Lineages: 7), Etoposide (Contrasts: 30; BioProjects: 7; Cell Lines: 25; Lineages: 8), and JQ1 (Contrasts: 20; BioProjects: 7; Cell Lines: 15; Lineages: 5).

We then performed fixed effects meta-analyses on the differential expression results across the selected contrasts for each compound. Meta-analysis on Decitabine contrasts resulted in 1,387 (9%) meta-up-regulated and 324 (2%) meta-down-regulated protein coding genes and 135 (15%) meta-up-regulated and 9 (1%) meta-down-regulated TE subfamilies (Figure 6A). Of the 135 up-regulated TE subfamilies, 88 (65%) were LTR ERVs. Doxorubicin, a DNA polymerase and topoisomerase inhibitor, showed 1,166 (8%) up- and 1,015 (7%) meta-down-regulated genes and 32 (3%) up- and 8 (1%) meta-down-regulated TE subfamilies. Of the 32 up-regulated TEs, 26 (81%) were from LTR ERVs. Etoposide, a topoisomerase inhibitor, showed 914 (7%) up- and 385 (3%) meta-down-regulated genes and 50 (6%) up- and 4 (*<*1%) meta-down-regulated TE subfamilies. Of the 50 up-regulated TEs, 42 (84%) were from LTR ERVs. JQ1, a bromodomain inhibitor, showed 1,162 (8%) up- and 1,891 (14%) meta-down-regulated genes and 38 (5%) up- and 18 (2%) meta-down-regulated TE subfamilies. Of the 38 up-regulated TEs, 31 (82%) were from LTR ERVs.

**Figure 6.**
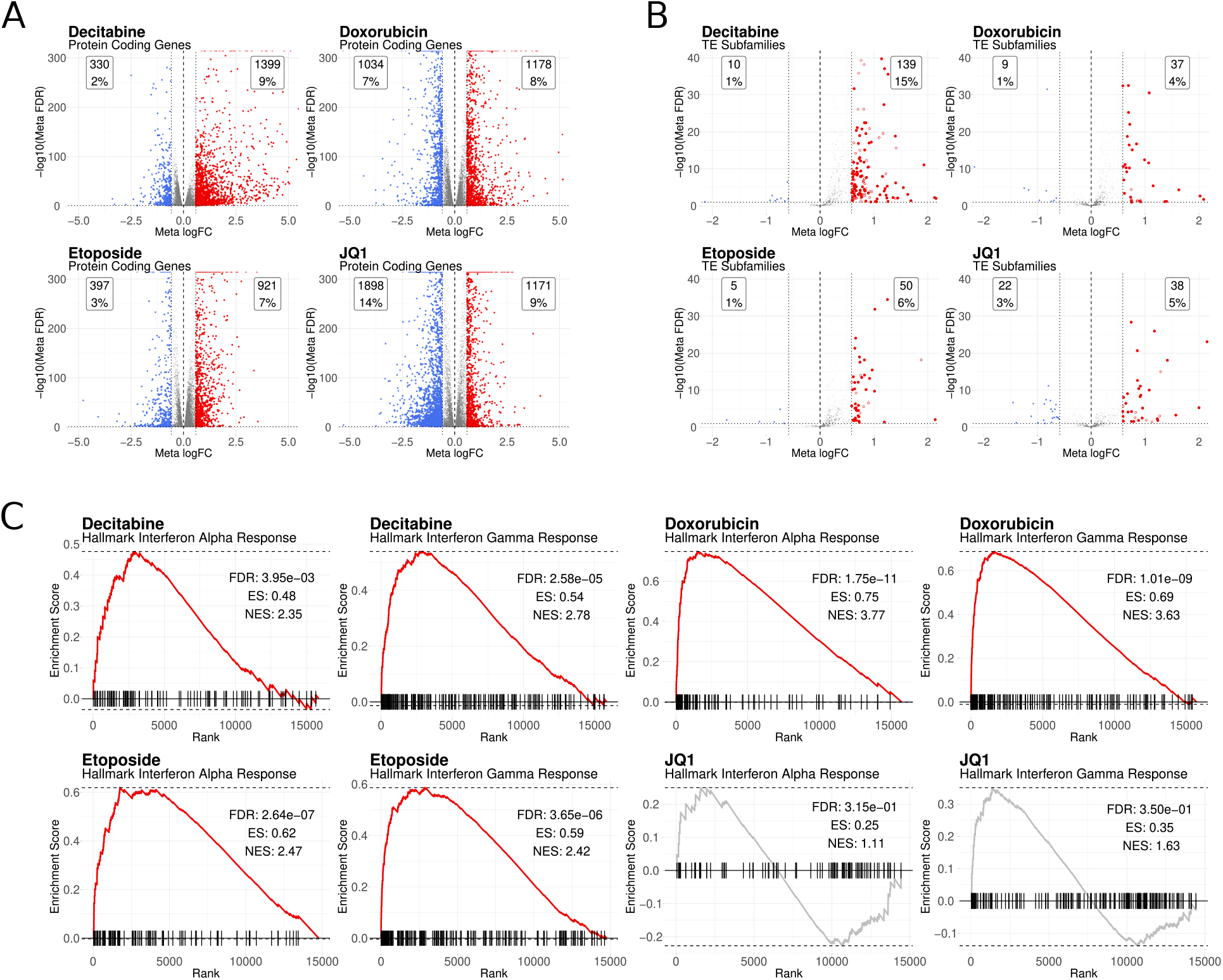
Meta-analysis of dysregulated protein coding genes and TE subfamilies. (A) Meta-volcano plots of protein-coding genes after performing fixed effect meta-analysis across all contrasts performed using the above compounds. Meta-volcano plots show the FDR-adjusted p-values and inverse-weighted logFC from fixed-effect meta-analysis of the gene or (B) TE subfamily data. Filled red points on the TE plots show any TE subfamily derived from an endogenous retroviral element (LTR ERV). (C) Competitive gene set testing results for each set of meta-analysis results. cameraPR() was used to assess enrichment using the meta-z-score as the ranking statistic. Enrichment was performed against the HALLMARK Interferon alpha and gamma response pathways.

Jackknife resampling was performed to assess the stability, bias, and variance of the differential expression estimates from each meta-analysis. Supplemental Figure 5 shows the jackknife estimated percentage differential expression, bias, and variance estimates. In all cases, the meta-analysis resulted in consistent and stable estimates of overall meta-differential expression. These results suggest the meta-analysis was able to discover consistently dysregulated features responding to specific compounds. To determine if these compounds were dysregulating similar gene/TE programs, an analysis of shared meta-significant features was performed. Protein coding genes had a larger number of shared up-regulated features as compared to TE subfamilies, which showed fewer shared meta-significant features (Supplemental Figure 6).

To test if the observed meta-up-regulation of ERVs was consistent with the induction of an innate immune response, we performed competitive gene set testing against the MSigDB [42] Hallmark Interferon Alpha Response and Hallmark Interferon Gamma Response gene sets on all genes ranked by their meta-z-statistic (Figure 6B). For all treatments except JQ1 we observed significant (FDR *<* 0.05) positive enrichment of both interferon alpha response and interferon gamma response. Taken together, the up-regulation of ERVs along with the induction of a consistent interferon response suggests that genotoxic compounds such as Etoposide and Doxorubicin also result in viral-mimicry-like signatures. It is important to note, however, that a more powerful mixed-effects meta-analysis may yield better control over the unexplored technical factors potentially influencing these results.

## Discussion

The role of transposable elements as immune modulators in cancer is an increasingly recognized area of research [43]. Traditional sequencing assays such as RNA-seq can capture the expression of TEs, however computational barriers, primarily due to multi-mapping of highly sequence-similar TE reads [3], result in TE specific counts being absent from large-scale re-processing efforts, limiting their study. Here, we sought to address this gap by uniformly reprocessing publicly available RNA-seq datasets to quantify TE and gene expression differences in cancer cell lines treated with epigenetic and cytotoxic compounds. Analysis of this reprocessed dataset confirmed the role of epigenetic compounds as inducers of viral mimicry responses. Interestingly, we also observed signatures of viral mimicry induction from several non-epigenetic compounds, such as CDK and topoisomerase inhibitors. Finally, to encourage reuse of this dataset, we developed a web application and database called TEDEdb which allows users to interactively explore and download all reprocessed data.

We observed considerable batch effects in the sample-level gene and TE count data. This result is unsurprising given the well-documented nature of batch effects with regard to transcriptomics profiling [31, 44, 45]. However, our analysis revealed that TEs are potentially more prone to batch effects than gene expression. An important caveat to this result is the confounding between each dataset and the cell lines studied. Many BioProjects present in our database performed experiments only on one or two cell lines, often of the same oncogenic lineage. This leads to confounding between experimental conditions and technical factors. For these reasons, we did not attempt to integrate the sample-level count data. While our results indicate important potential technical differences observed when quantifying and analyzing TEs, further analysis is needed to determine the causes of these differences.

To mitigate batch effects and extract meaningful patterns of TE and gene differential expression across disparate treatments and cell lines, we instead focused on comparing differential expression results across studies. We observed that most perturbations did not result in the dysregulation of many genes or TE subfamilies. However, a small minority of contrasts showed a high degree of up-regulation of TE subfamilies. Within this subset of contrasts we observed many DNMTi treated samples, which have been previously shown to up-regulate both silenced genes [46] and endogenous retroelements [47]. Interestingly, we also observed that some compounds, not traditionally regarded as epigenetic drugs, showed a large degree of TE induction. CDKis and topoisomerase inhibitors were among the top mechanisms resulting in the induction of TEs. Mechanistically, these results are consistent with previous studies and emerging literature which suggests targeting CDKs can cause the epigenetic re-activation of silenced genes and the relaxing of chromatin leading to the expression of TEs [28, 29, 48]. Likewise, targeting topoisomerases is a well-established anticancer treatment resulting in the de-repression of genes and TEs [49, 50].

We used meta-analyses to combine differential expression results from diverse cell lines and treatments to find common signatures of dysregulated genes and TEs, and to test if these signatures align with a viral mimicry phenotype. We observed that Decitabine treated samples showed the highest degree of TE up-regulation along with an enrichment of interferon alpha and gamma response genes. Etoposide, Doxorubicin, and JQ1 had a smaller magnitude of TEs induced. However, of the TEs induced by these treatments, most were endogenous retroelements. The induction of ERVs by JQ1, a bromodomain inhibitor, is generally less well studied [51]. JQ1 has been observed to induce LTR expression in H1299 cells [52]. While this analysis provides compelling evidence of conserved TE dysregulation signatures consistent with previously reported findings, a more powerful meta-analysis using a random effects model is potentially more suitable. Such models may be required for extracting robust conserved signatures of viral mimicry response induced by non-canonical epigenetic compounds.

Finally, we developed a user-friendly web application that can be used to access the processed data and perform analyses on the differential expression results. The web application is publicly available at: https://dataexplorer.coriell.org/TEDEdb/. This web application allows users to select among the more than 2,000 differential expression results and perform dimensionality reduction and meta-analysis on selected datasets. Users can explore and download differential expression results for any contrast in the database. Gene set enrichment analysis, gene ontology over-representation testing, and visualization of gene and TE differential expression can also be performed on the platform. To encourage re-use of this data and to enable researchers to perform their own analysis, we have also made all of the ∼51TB of raw count data available for download. We hope that this database and resource encourages the exploration of TEs in the context of cancer treatment, moving beyond canonical epigenetic inducers of TEs and towards the discovery of new mechanisms of induction and treatment response.

## Materials and Methods

### Data collection

Experimental samples were queried using the metaSRA web interface [53]. Human cancer cell lines or primary samples from RNA-seq studies treated with cytotoxic or epigenetic therapies were selected for inclusion. To ensure robust differential expression estimates, studies must have included control and treatment conditions, with at least 3 biological replicates per group. Treatment conditions, in this context, are defined as a cancer cell line or primary patient sample, treated with an epigenetic or cytotoxic drug.

Run and BioSample information, for each sample, was downloaded from the SRA and parsed into a machine-readable format. Because NCBI doesn’t require standardized metadata for SRA submissions, we manually added annotations to samples using the provided BioSample information and the original publication (if available). Such manual annotation of each sample included extracting and annotating cell lines with standardized names derived from Cellosaurus [37] and The Cancer Cell Line Encyclopedia [38], creating new labels for the tissue-of-origin, and disease state for each cell type, extracting and annotating drug-treated samples with standardized drug names and targets from The Broad Drug Repurposing Hub [40] or PubChem, and creating new group labels for downstream differential expression analysis. Once manual annotation was complete for each sample, raw fastq files were downloaded using the sra-tools prefetch and fasterq-dump tools, respectively, before being input into the quantification step.

### Quantification

We then quantified raw fastq files against a modified version of the REdiscoverTE transcriptome index [21], using Salmon [24]. The REdiscoverTE transcriptome consists of distinct transcript sequences from GENCODE v26 basic, all RepeatMasker elements from standard chromosomes, and distinct sequences representing GENCODE TE-containing introns [21]. Since the publication of the original REdiscoverTE method, Salmon developers showed that more accurate transcript quantification could be achieved by augmenting the transcriptome index with genomic decoys [54]. Therefore, we included a full genomic decoy of the GENCODE v26 primary fasta within our version of the REdiscoverTE transcriptome index.

The transcript and TE-locus level counts were imported into R using the tximport package, which incorporates transcript-level information into gene-level abundance estimates and TE loci-level information into subfamily-level counts, improving downstream differential expression analysis [55]. This is another modification to the original REdiscoverTE pipeline. After raw counts were imported, for each run, mapping statistics were extracted from the Salmon logs and used to flag potential outlier samples. A sequencing run was flagged as a potential outlier if any of the following mapping statistics exceeded 3 Median Absolute Deviations (MAD) from the median: fragment length, number of equivalence classes, number of processed reads, number of mapped reads, number of decoy fragments, number of dovetail alignments, or number of filtered fragments.

### Differential Expression Analysis

Tximport [55] was used to import quantification results and summarize transcript-level counts to the gene-level and TE-locus level counts to the subfamily level. TE subfamily-level counts were chosen for downstream analysis for ease of interpretability and comparison to previously published results. Feature count matrices were filtered for features with *>* 10 counts-per-million in 70% of samples, and normalized. Due to the nature of processing of a large number of samples in a semi-automated fashion, as well as incomplete experimental information from GEO or the original publication, we could not assume that count distributions within a given experiment conformed to the assumptions of global scaling normalization methods like Trimmed Mean of M-values (TMM) [56] or DESeq2’s median of ratios [57]. Even though some departures from the assumptions of these methods can be tolerated, large global expression shifts, often observed in drug-treated cancer RNA-seq studies, can lead to spurious downstream differential expression results [58, 59]. We therefore employed a data-based method implemented in the quantro [59] R package to estimate the magnitude of global expression shifts in the raw count data to inform normalization. For experiments where global expression assumptions were found to be violated, either by having a permutation-based quantro F-statistic empirical p-value *<* 0.01, or an observed difference in group medians (p *<* 0.01 by ANOVA), smooth-quantile normalization was used from the qsmooth R package [58]. Smooth-quantile normalization assumes that the statistical distribution of counts, for each sample in a group, has the same distributional shape, allowing for global distributional differences between groups. This normalization method allows for large shifts in global expression, commonly observed in drug-treated cancer samples, to be preserved, while also accounting for technical artifacts, but with the drawback of making strong assumptions about the nature of these shifts. If the experiment was found not to violate the assumptions of global scaling normalization, TMM normalization factors were used in the downstream analysis, as calculated in edgeR [60].

Following estimation of normalization factors, qsmooth- or TMM-normalized log2-counts-per-million values were used as input for differential expression testing with limma [61], using the treat function. This function tests differential expression between conditions, relative to a fold-change, improving false discovery rate (FDR) estimation [62]. Experimental contrasts used in differential expression testing for each experiment were manually annotated based on the available metadata information, to the best of our ability. Absent available code from the original publication, it is possible that experimental contrasts differed from those used by the authors.

For easier sharing and reproducibility of analyses, for each experiment, a SummarizedExperiment object was saved containing the original and normalized count matrices, group-level metadata, the fitted model object, and differential expression testing results for all contrasts. The resulting SummarizedExperiment objects and raw counts are available in the database.

### Web-Application Utilities

The primary utility of our TEDEdb database can be most effectively appreciated through the use of the accompanying Shiny application, which provides a simple yet comprehensive interface for performing meta-analysis of differential expression results or exploratory analysis of individual differential expression results. The app is divided into several pages, as described below.

The meta-analysis page enables exploration of DE results across many experiments. Each of its subpages allows for different levels of analysis across aggregated differential expression results. The first of the sub-pages is the “Data Selection” page, which provides an interface for selecting contrasts (e.g., Treatment vs Control comparisons) in all subsequent “meta-analysis” pages. Users can use the dropdowns to dynamically select subsets of contrasts by their BioProject ID, drug class, tissue type, etc., to hone in on specific dysregulation patterns that may, for example, be present following a given drug treatment.

Following data selection, PCA or UMAP dimensionality analysis can be performed on the selected contrasts. Here, the user can perform dimensionality reduction (PCA or UMAP) on genes alone, TEs alone, or both. The input data values for dimensionality reduction are one of the values computed from the differential expression analysis for a particular feature, e.g., logFC, P-value, or z-statistic. Using these values for dimensionality reduction allows for exploring differential expression patterns across disparate treatments, cell lines, experiments, etc. After performing dimensionality reduction, a scatter plot of the computed embeddings, or PCs, is displayed to the user. Users can color points using the available metadata or select subsets of points for further inspection. Selected points are displayed in a table below the plots, along with accompanying metadata.

In addition to visualizing common dysregulation patterns, users can also directly perform meta-analysis, using one of the built-in p-value combination methods. After computing a p-value combination for all features of the selected contrasts, a meta-volcano plot is displayed, showing the combined p-values and representative logFC values. These plots allow users to quickly examine what genes or TEs are commonly and significantly dysregulated across many differential expression experiments.

Users can also probe the differential expression results for individual experiments, using the differential expression page, which displays interactive volcano and MA plots and a table of the given DE results. Users can adjust threshold values on-the-fly and separately examine TEs or genes. In addition to viewing differential expression results, users can perform gene set enrichment or over-representation analysis for any of the selected contrasts, using the “GSEA” and “Over-representation” tabs, respectively.

### UMAP Dimensionality Reduction of Sample-level Counts

UMAP dimensionality reduction was performed on principal components computed from TMM normalized log2 counts-per-million values of genes and TEs for each BioSample after removing those that were flagged as outliers based on mapping statistics. PCA was performed such that the number of PCs chosen for input into UMAP represented ∼80% of the total variation of the original dataset for both gene and TE expression matrices.

PERMANOVA analysis was performed on the Euclidean distance matrices computed on the PCs above using the adonis2 [63] function from the vegan R package [64] treating oncotype lineage and BioProject sequentially as predictor variables for both gene and TE matrices.

### Meta-analysis and Competitive Gene Set Testing

Meta-analyses were implemented using an inverse-variance weighted fixed-effects model. We chose this approach to prioritize computational efficiency and real-time user interactivity during large-scale data exploration within the app. We consider these results to be high-level signatures intended for hypothesis generation. To facilitate more specialized downstream analyses, including mixed-effects or random-effects modeling tailored to specific biological questions, all processed count data and associated metadata are provided for public download.

Meta-analysis was performed on contrasts from Decitabine, Doxorubicin, Etoposide, and JQ1 treated samples after excluding any contrasts from primary cells or multiple cell lines, any non-cancerous cell lines, or any contrasts in which a sample was flagged as a sequencing outlier. For each set of contrasts, for each gene/TE subfamily, a meta log2 fold-change value was computed using inverse-variance weighting on the standard errors computed from limma::topTreat. These meta log2 fold-change values were used to derive a meta z-score and meta p-value for each gene within the set of contrasts. Meta z-scores were used as a ranking statistic to perform competitive gene set testing with CAMERA [65] on the MSigDB hallmark interferon alpha response and hallmark interferon gamma response gene sets. Jackknife (leave-one-out) resampling was performed on each set of contrasts to assess stability, bias, and variance of the estimated percentage of differentially expressed features. To create background estimates of meta-differential expression, feature-level (genes/TEs) statistics were randomly permuted within each contrast and meta-analysis was performed on the permuted data. This process was repeated 500 times. 95% CIs were computed around the mean percentage meta-dysregulated features across all permutations.

## Data and Code Availability

All raw fastq files are publicly available for download on NCBI SRA. The web application can be accessed at: https://dataexplorer.coriell.org/TEDEdb/. All Shiny application codes are freely available on GitHub: https://github.com/coriell-research/TEDEdb-app. Individual analysis scripts for all analyzed experiments, as well as the pipeline scripts for downloading, processing, and data preparation, are publicly available on GitHub: https://github.com/coriell-research/TEDEdb.

## Figures

**Supplemental Figure 1.**
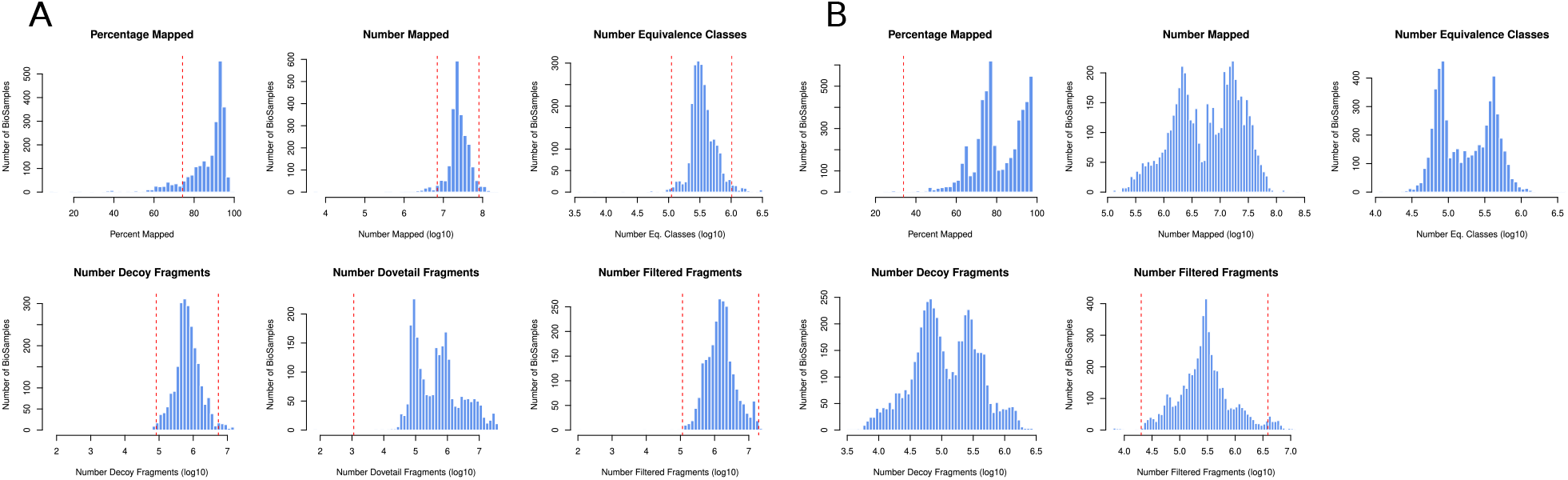
Distribution of Salmon mapping statistics for all BioSamples included in TEDEdb. (A) Paired-end sequencing samples. From left to right, Percentage of mapped fragments, absolute number of mapped fragments, the number of equivalence classes detected, number of fragments aligned to the decoy sequences, number of fragments with dovetail alignments, the number of alignments filtered due to mapping quality. +/-3 Median absolute deviations (MAD) are shown as the dotted red line. Samples outside of this fence are flagged as potential outliers. (B) Single-end sequencing samples.

**Supplemental Figure 2.**
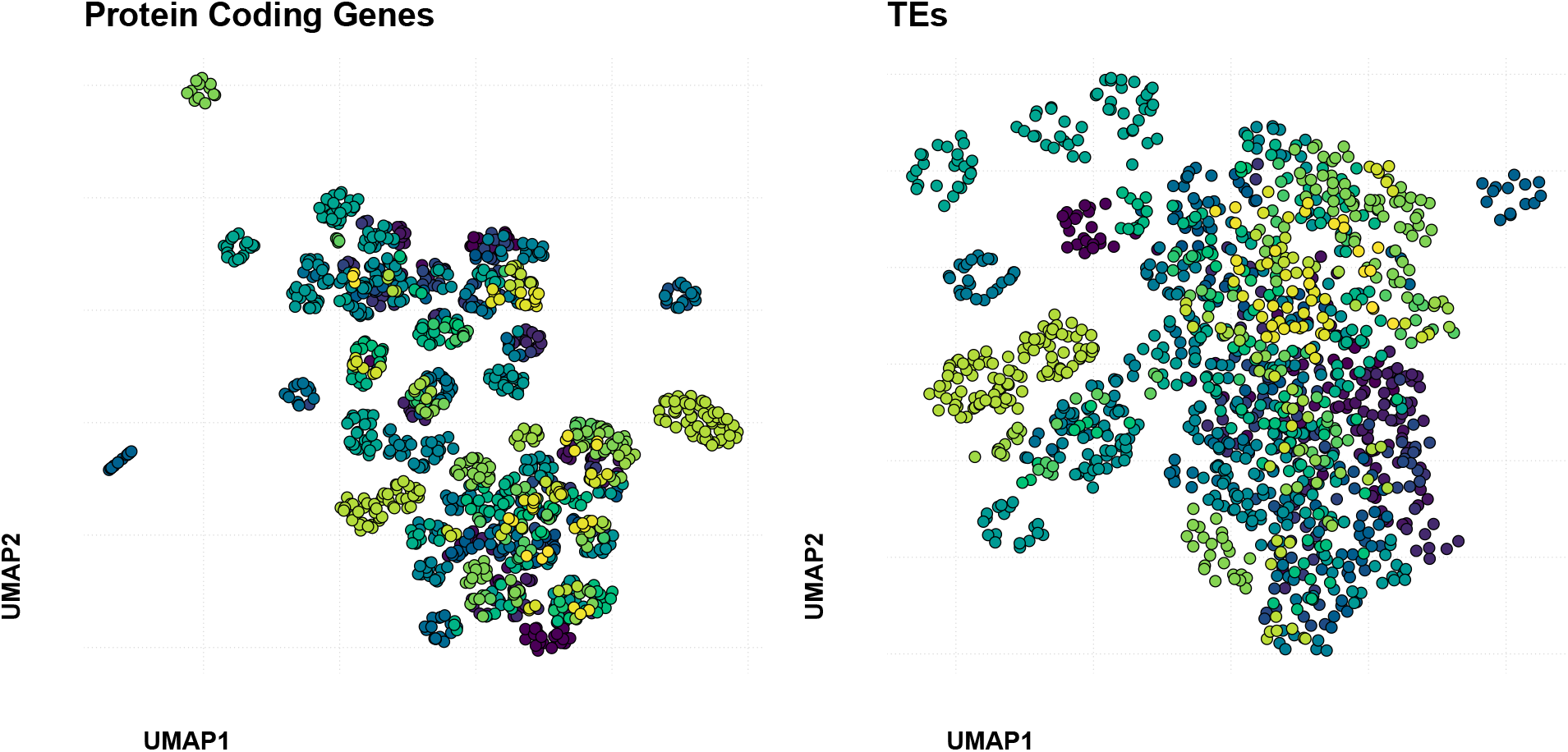
UMAP reduction of BioSamples by gene and TE expression. UMAP dimensionality reduction on protein coding gene expression (left) and TE subfamily expression (right) performed on control BioSamples present in TEDEdb. Samples have been colored by their BioProject (n=138) of origin. The smallest subclusters in both UMAP plots indicate substantial technical artifacts attributable to the given samples’ BioProject. Larger clusters, as shown in Figure 2, tend to capture cell lineage information which tends to be preserved in both gene and TE UMAPs.

**Supplemental Figure 3.**
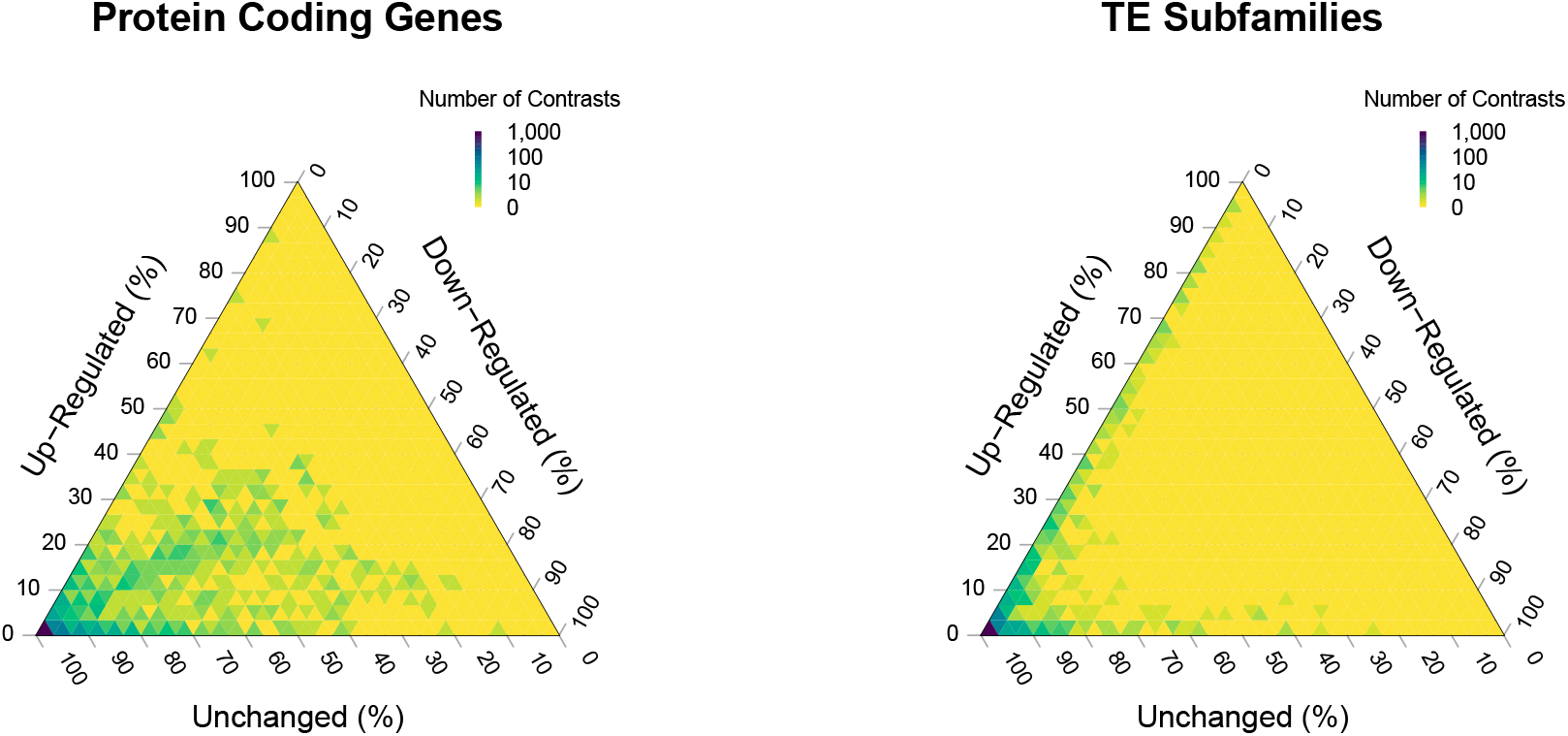
Ternary plot showing percentage of dysregulated features across contrasts in the TEDEdb. The majority of contrasts do not show many differentially expressed genes or TEs. (Left) Differential gene expression tends to be along the diagonal indicating even proportions of up- and down-regulated genes. In contrast, TE subfamilies are mostly unaffected. However, a subset of contrasts shows a striking degree of up-regulation of TE subfamilies.

**Supplemental Figure 4.**
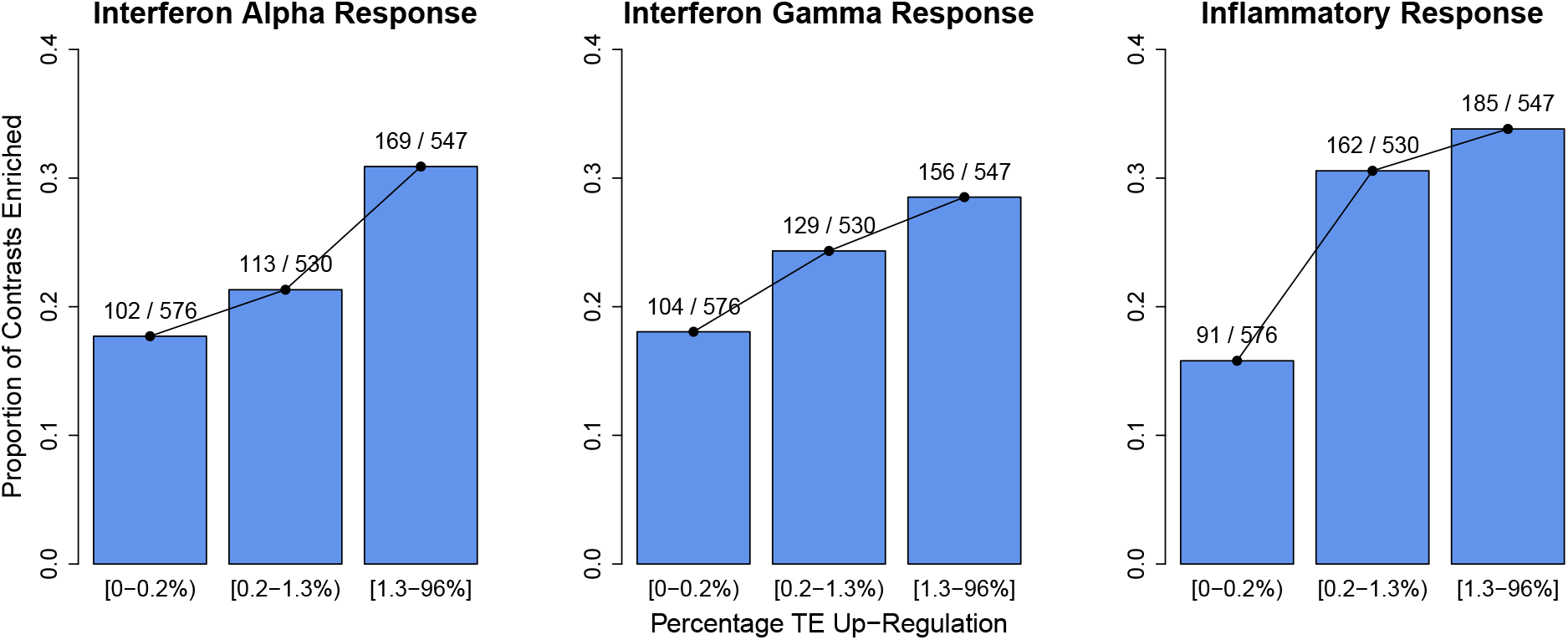
Proportion of contrasts by TE up-regulation and enrichment of viral mimicry signatures. Bars show the proportion of contrasts by TE expression with the number of contrasts listed above (enriched / enriched + not enriched), dichotomized by percentage of TE up-regulation by tertile. Facets stratify the data by HALLMARK pathways.

**Supplemental Figure 5.**
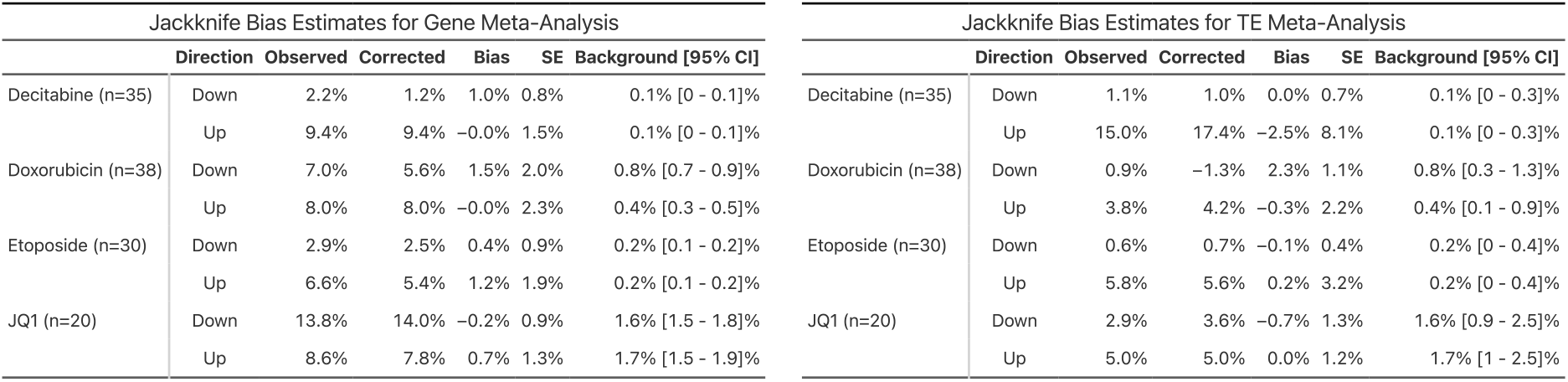
Jackknife bias estimates on gene and TE meta-analysis. Rows show the compound and number of contrasts included in the meta-analysis. Direction indicates the direction of the differential expression. Observed (%) represents the percentage of differentially expressed features in the full meta-analysis. Corrected (%) represents the Jackknife bias-corrected percentage of differential expression. Bias and SE represent the bias and standard error of the Jackknife estimator.

**Supplemental Figure 6.**
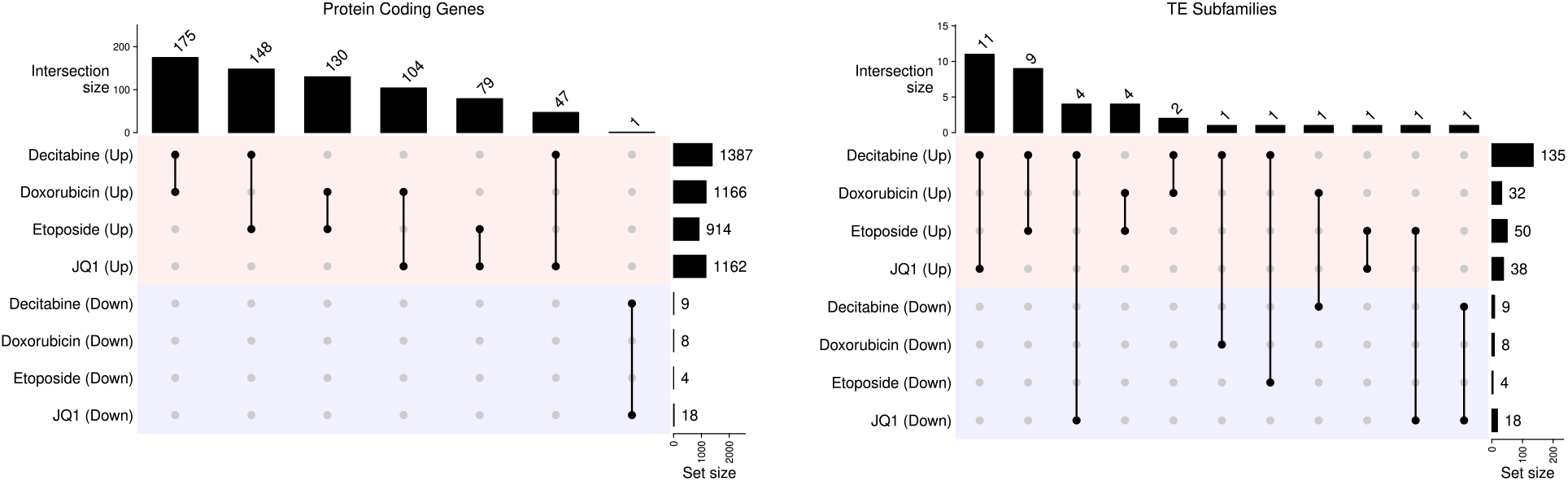
Shared meta-significant genes and TE subfamilies across selected compounds. UpSet plots showing the number of shared protein coding genes and TE subfamilies for each set of pairwise comparisons.

## Notes

### Competing Interest Statement

The authors have declared no competing interest.

